# Inhibitory signaling in mammalian olfactory transduction potentially mediated by Gα_o_

**DOI:** 10.1101/811125

**Authors:** Elizabeth A. Corey, Kirill Ukhanov, Yuriy V. Bobkov, Jeremy C. McIntyre, Jeffrey R. Martens, Barry W. Ache

## Abstract

Olfactory GPCRs (ORs) in mammalian olfactory receptor neurons (ORNs) mediate excitation through the Gα_s_ family member Gα_olf_. Here we tentatively associate a second G protein, Gα_o_, with inhibitory signalling in mammalian olfactory transduction by first showing that odor evoked phosphoinositide 3-kinase (PI3K)-dependent inhibition of signal transduction is absent in the native ORNs of mice carrying a conditional OMP-Cre based knockout of Gα_o_. We then identify an OR from native rat ORNs that are activated by octanol through cyclic nucleotide signaling and inhibited by citral in a PI3K-dependent manner. We show that the OR activates cyclic nucleotide signaling and PI3K signaling in a manner that reflects its functionality in native ORNs. Our findings lay the groundwork to explore the interesting possibility that ORs can interact with two different G proteins in a functionally identified, ligand-dependent manner to mediate opponent signaling in mature mammalian ORNs.

## Introduction

ORs comprise the largest family of mammalian GPCRs (Buck and Axel, 1991). Ligand (odorant) binding to ORs results in the cyclic nucleotide-dependent excitation of ORNs through Gα_olf_, a member of the Gα_s_ subfamily (e.g., Belluscio et al., 1998). It has been known for some time that olfactory perception shows ‘mixture suppression’ and ‘mixture synergism’, in which one odorant either reduces or enhances, respectively, the percept of another (e.g., Cain, 1974; Laing et al., 1984), and that at least some of this perceptual modulation can be assigned to the olfactory periphery (e.g., Bell et al., 1987; Laing and Wilcox, 1987). Receptor-driven modulation has since been studied directly (see following paragraph) and was recently shown to be widespread across ORs, indicating that it makes a fundamental contribution to the peripheral olfactory code (Xu et al., 2019). Thus, it is important to understand the processes that modulate cyclic nucleotide-dependent excitation in the dynamic range of activation.

Receptor-driven ‘mixture suppression’, also referred to as inhibition, antagonism, or masking, has received the most attention. Pharmacological, physiological, and computational evidence ascribe odor-evoked inhibition to competitive antagonism (e.g., Firestein and Shepherd, 1992; Kurahashi et al., 1994; Oka et al., 2004). The implication is that ‘mixture suppression’ results from a reduction in cyclic nucleotide-dependent excitation due to odorants competing for a common binding site on the OR. Both physiological (Rospars et al, 2008) and computational (Reddy et al., 2017) evidence ascribe odorant-evoked inhibition to non-competitive antagonism in addition to competitive antagonism. Multiple non-competitive processes can result in odorant-evoked inhibition. Some, such as ‘odor masking’ involving the non-specific action of the antagonist on the cyclic nucleotide gated (CNG) output channel (e.g., Takeuchi et al., 2009), cannot account for the broad ligand specificity of odor-evoked inhibition seen across ORNs (Xu et al., 2019). A non-competitive process linked to odorant-evoked inhibition that is consistent with the ligand specificity seen across ORNs involves phosphoinositide 3-kinase (PI3K)-dependent signaling (Spehr et al., 2002; Ukhanov et al., 2010, 2011, 2013; Yu et al., 2014). Interestingly, the primary product of PI3K-dependent signaling *in vivo*, PtdIns (3,4,5)P3 (PIP3), competitively competes with cAMP-dependent activation of the CNG channel (Zhainazarov et al., 2004; Brady et al., 2006), potentially confounding a simple mechanistic understanding of receptor-driven ‘mixture suppression’. Pharmacological evidence that PI3K-dependent, odorant-evoked inhibition is mediated by a Gβγ subunit implicates a G protein complex in this process (Ukhanov et al., 2011), as does earlier evidence that in heterologous systems at least, the function of an odorant (agonist, antagonist) depends on the G protein used (Shirokova et al., 2005).

Implicating a Gβγ subunit in PI3K-dependent, odorant-evoked inhibition raises the question of the associated Gα protein. While Gα_olf_, the most abundant Gα isoform expressed in the cilia of mammalian ORNs, could mediate activation of PI3K signaling through the release of Gβγ, other isoforms occur in cilia-enriched membrane preparations from the olfactory epithelium (OE) (e.g., Schandar et al., 1998; Wekesa and Anholt, 1999; Mayer et al., 2009). These other G proteins may function in processes as diverse as adaptation and cell survival (Watt et al., 2004; Mashukova et al., 2006; Kim et al., 2015a,b), but have also been implicated in signal transduction (e.g., Scholz et al., 2016b). If two different G protein complexes are involved in olfactory signal transduction, it is important to understand whether both are activated by the same OR.

Here, we provide evidence potentially linking PI3K-mediated inhibitory signaling pathway to Gα_o_. We demonstrate that odor-evoked PI3K-dependent inhibitory signaling is no longer detectable in mice carrying an OMP-Cre conditional deletion of Gα_o_. We show that fluorescently-tagged Gα_o_ is trafficked to the cilia of native ORNs using viral-mediated ectopic expression, and that Gα_o_ expression is reduced in the ORNs of mice carrying the OMP-Cre conditional deletion of Gα_o_ using IHC. We then use single cell RT-PCR to identify an OR expressed by mammalian ORNs that were activated by octanol and inhibited by citral in a PI3K-dependent manner. The functionality of the identified OR (Olr1845) persists in a HEK293T-based pCRE-SEAP assay. Using the same expression system we then implicated Gα_o_ in odor-dependent activation of PI3K by that OR using an ELISA. Collectively, our results are consistent with, although do not prove, that mammalian ORs can interact with at least two different G proteins in a functionally identified, ligand-dependent manner.

## Methods

### Animals

Experiments were performed on adult female Sprague-Dawley rats, adult CD1 mice, adult M71-SR1-IRES-tauGFP mice, as well as adult C57BL/6 and *cGnαo^-/-^* mice. All animal procedures were performed in accordance with the University of Florida animal care committee’s regulations. Animals were euthanized by inhalation of carbon dioxide and decapitated immediately prior to dissection. All experiments were performed at room temperature (22–25°C) unless otherwise noted. *cGnαo^-/-^*animal breeding, genotyping, and genomic DNA analyses were performed using published protocols and primers (Chamero et al., 2011; Choi et al., 2014).

### In situ hybridization and immunolabeling of cryosections

Tissue fixation and cryo-sectioning were performed using published protocols. Briefly, the OE was fixed in 4% paraformaldehyde and then the tissue soaked in 30% sucrose at 4°C before embedding in optimal cutting temperature medium. 12 μM sections were collected under RNase-free conditions and stored at −80°C until use. *In situ* hybridization was performed using a modification of published methods (Ishii et al., 2004; Choi et al., 2016a). Briefly, tissue sections were hybridized with digoxigenin-labeled riboprobes for *Gnao* and *OMP* detection. After washing to remove unbound probe, the sections were then incubated with anti-digoigenin-HRP antibody (Roche) and labeling was detected with NBT/BCIP (Sigma). The sections were cover-slipped with Fluormount with DAPI (Southern Biotechnology) and visualized with a 10x and an oil immersion 60x lens on an Olympus BX41 microscope.

Immunostaining was performed using modifications of published protocols (e.g., Choi et al., 2016). Briefly, antigen retrieval was performed by incubating the slides with 10 mM sodium citrate buffer (pH 6.0) at 60°C for 30 min. After blocking with 10% (vol/vol) normal goat serum, 1% BSA, and 0.1% Triton X-100, sections were incubated with primary antibodies overnight at 4°C. The antibodies included Gα_o_ (rabbit, 1:200; Santa Cruz Biotechnology) and OMP (goat, 1:500; Wako). The slides were washed with PBS containing 0.1% Triton X-100 and then with secondary antibodies conjugated with Alexa Fluor 488 (Invitrogen). Slides were coverslipped with Fluoromount DAPI (Southern Biotech) and labeling was visualized with 10x and oil immersion 60x lenses.

### Calcium imaging

Acutely dissociated rat or mouse ORNs were imaged using standard published approaches. Briefly, olfactory epithelia were dissected in ice-cold modified artificial cerebrospinal fluid (ACSF) saturated with 95% O_2_ and 5% CO_2_ that contained (in mM): 120 NaCl, 25 NaHCO_3_, 5 KCl, 1.25 Na_2_HPO_4_, 1 MgSO_4_, 1 CaCl_2_, 10 glucose. The tissue was transferred in low-Ca^2+^ (0.6 µM free Ca^2+^ buffered with 5 mM EGTA) ACSF supplemented with 0.5 mg/ml papain (Sigma-Aldrich) and, in some cases, 10 units/ml TurboDNAse (Promega). After incubation for 20 min at 37°C in 5% CO_2_, the tissue was gently washed with normal oxygenated ACSF several times, minced with a razor blade and triturated with a large bore fire polished glass pipette. The resulting suspension was filtered through a 40 µm cell strainer (BD BioSciences). An aliquot of the suspension was mixed with 10 µM Fluo-3 or Fluo-4 containing 0.04% Pluronic F127 and placed on a glass coverslip coated with concanavalin A (Sigma-Aldrich) in a recording chamber (RC22, Warner Instruments). The volume of the chamber was 200 µL, allowing for complete exchange of the solution during application of odorant and/or inhibitors. In some experiments cells were placed and imaged in 35mm tissue culture dishes with cover glass bottom (FluoroDish, WPI) treated with concanavalin A. Odors were applied using a multi-channel rapid solution changer (RSC-160, Bio-Logic). The cells were illuminated at 500 nm and the emitted light was collected at 530 nm by a 12-bit cooled CCD camera (ORCA-R2, Hamamatsu). Both the illumination and image acquisition were controlled by Imaging Workbench 6.0 software (INDEC BioSystems). Each cell was assigned a region of interest (ROI) and changes in fluorescence intensity within each ROI were analyzed. Continuous traces of multiple responses were compensated for slow drift of the baseline fluorescence when necessary. All recordings were performed at room temperature (22-25°C). Single odorants were of highest purity obtained from Sigma-Aldrich and were prepared fresh as used from 0.5M DMSO stocks kept at −20°C. The complex odorant Henkel-100 was dissolved 1:1 in anhydrous DMSO as a working stock solution.

### Viral expression of fluorescently tagged Gα_o_

GFP and mCherry were inserted into the coding sequence of mouse Gα_o_ using site directed mutagenesis to create EcoRI cut sites within the Gα_o_ coding sequence followed by restriction enzyme digestion and T4 ligation. GFP and mCherry were amplified by PCR with primers designed to allow in frame insertion as previously described (Hynes et al., 2004). All constructs were fully sequenced prior to use. Gα_o_:GFP adenovirus (AdV) and Gα_o_:mCherry adeno-associated virus (AAV2/5) were produced using previously described methods (e.g., Zolotukhin et al., 2002; McIntyre et al., 2015). For expression using AV in native tissue, recombinant GFP-fused cDNA was cloned into the vector p-ENTR by TOPO cloning methods. The inserts were then recombined into the adenoviral vector pAD/V5/-dest using LR Recombinase II (Life Technologies, Carlsbad CA). Viral plasmids were digested with PacI and transfected into HEK293 cells. Following an initial amplification, a crude viral lysate was produced, and used to infect confluent 60-mm dishes of HEK293 cells for amplification according to the ViraPower protocol (Life Technologies). AdV was isolated with the Virapur Adenovirus mini purification Virakit (Virapur, San Diego, CA), dialyzed in 2.5% glycerol, 25 mM NaCl and 20 mM Tris-HCl, pH 8.0, and stored at −80°C until use. For ectopic expression in native tissue using AAV, the Gα_o_:mCherry fusion was cloned into the pTR-UF50-BC plasmid vector and virus was propagated in HEK293 cells using the pXYZ5 helper plasmid. For viral transduction of ORNs, mice were anesthetized with a Ketamine/Xylazine mixture and 10-15 µL of purified viral solution was delivered intranasally as a single injection per nostril. Animals were used for experiments at 10 days post-infection. The entire turbinate and septum were dissected and kept on ice in a petri dish filled with oxygenated ACSF. For imaging a small piece of the OE was mounted on the stage of the microscope in a perfusion chamber with the apical surface facing down. High resolution *en face* imaging of freshly dissected OE was performed on an inverted confocal microscope Leica SP5. Images were processed using ImageJ (NIH http://imagej.nih.gov/ij/) and assembled in CorelDraw13 (Corel).

### Single Cell RT-PCR

Rat ORNs functionally characterized by calcium imaging were collected with a sterile glass micropipette directly into RT buffer for lysis. Cells were immediately frozen at stored at −80°C. Single cell RT-PCR was performed using a modified approach based on previously described methodology (Touhara et al., 1999). Briefly RT was performed using a Verso RT kit (Thermo Fisher) with an anchored oligo dT primer for 60 minutes at 42°C. RT was followed by PCR detection of OMP and beta actin to exclude cells that were not ORNs and samples contaminated with genomic DNA. PCR with degenerate primers designed to amplify OR genes was performed as follows. The first round of amplification of OR genes was performed in a solution containing 0.4 μM each of the published degenerate primer and an adapter primer targeting the oligo d(T)18-anchor used for the RT, 0.2 mM dNTP, and PrimeSTAR HS Taq (Clontech) and the second amplification used a nested set of primers targeting ORs. Each PCR consisted of 5 min at 95°C followed by 40 cycles at 95°C for 1 min, an annealing temperature dependent on primers for 3 min, and 72°C for 2 min. The PCR products were subsequently cloned into pGEM-T Easy (Promega) followed by sequencing (McLab) of multiple clones for each PCR product.

### OR expression constructs

Rat ORs identified by single cell RT-PCR were amplified from genomic rat DNA and mOR261-1 was amplified from genomic mouse DNA. The ORs were cloned into a pME18S-based Lucy-Rho vector (denoted here as pLucy-Rho-OR) (Shepard et al., 2013) for mammalian expression. All constructs were sequenced prior to use.

### Culture and transfection of HEK293T cells

HEK293T cells (ATCC) were grown in Dulbecco’s modified Eagle medium (DMEM) supplemented with 10% fetal bovine serum, penicillin (100 U/ml), and streptomycin (100 mg/ml) in 5% CO_2_ at 37°C. Before transfection, the cells were seeded into 35 mm tissue culture treated dishes and incubated for 24 hours. For pCRE-SEAP and PI3K assays, cells were transfected at 70% confluency using X-treme-GENE HP (Roche) at a ratio of 3:1 with plasmid DNA following the manufacturer’s instructions.

### pCRE-SEAP assay

cAMP production was measured as previously described (Durocher et al., 2000). HEK293T cells were transfected with the expression vectors pcDNA3.1 Ric-8b (50 ng; generously provided by Dr. Bettina Malnic, Universidade de São Paulo, Brazil), pcDNA3.1(+) Gαolf (50 ng; Missouri S&T cDNA Resource Center), pcDNA3.1(+) RTP1s (100 ng; subcloned from construct purchased from Thermo Fisher) and pLucy-Rho-OR (1.5 μg). For control experiments cells were transfected as above, however, the pLucy-Rho-OR construct was omitted. Cells were also transfected with 1.5 μg of a pCRE-SEAP, where the expression of the secreted alkaline phosphatase (SEAP) is under regulation of the cAMP responsive elements, (pCRE-SEAP) or a pTAL-SEAP, where the cAMP responsive elements are not present (Clontech; Durocher et al., 2000). Cells were also transfected with 50 ng pcDNA5/TO/LACZ (Invitrogen) to assess transfection efficiency. At 24 hr post-transfection the cells were re-seeded for SEAP analysis and odorants were added at the indicated dilutions at 48 hours post-transfection. Cells and supernatants were collected 20 hr later and centrifuged for 5 min at 5000 g. The supernatants were incubated for 30 min at 65°C and then frozen until analysis. SEAP activity was measured by mixing 100 μl of supernatant with an equal amount of BluePhos substrate (KPL). Samples were monitored for color development at 630 nm in a microwell plate reader. Mean SEAP activity was determined after subtracting the response of cells that were not expressing an OR and is reported in OD630 arbitrary units +/- SEM. Each experiment was repeated in triplicate with three replicates each. The data were analyzed using GraphPad Prism.

### PI3K assay

HEK293T cells were transfected with 1.5 μg of pME18s Lucy-Rho-OR, 100 ng of pcDNA3.1(+) RTP1s and 100 ng of the indicated G protein construct, as well as with 0.5 μg of pBTK-PH-YFP (generous gift from Dr. Tamas Balla; (Balla et al., 2009). pcDNA3.1(+)-based constructs for Gα_o_, Gα_olf_, and Gα_oG203T_ were obtained from the Missouri S&T cDNA Resource Center. At 24 hours post-transfection, cells were split into 35 mm dishes for analysis. After 24 hours, cells were incubated for 1 hour in 0.5% fetal bovine serum in DMEM including phosphatase inhibitors (Boston Bioproducts). For PI3K activation, cells were treated with odorant or DMSO (odorant carrier) for 30 sec and then immediately lysed with ice cold 5% TCA. Cells were scraped from dishes and the lysates were stored immediately at −80°C until analysis. For analysis, lipids were extracted following a chloroform:methanol protocol and used immediately in a PIP3 ELISA following the manufacturer’s instructions (Echelon Biosciences). Each experiment was performed in triplicate. Mean PIP3 production was determined by subtracting the response to DMSO and is presented as ΔPIP3 (pM). The data were analyzed using GraphPad Prism.

## Results

### Gα_o_ is required for PI3K-dependent inhibitory signal transduction in mouse ORNs

Given varied lines of evidence that Gα_o_ is expressed in mammalian ORNs (Mayer et al., 2009; Keydar et al., 2013; Heron et al., 2013; Nickell et al., 2012; Omura and Mombaerts, 2014; Saraiva et al., 2015; Scholz et al., 2016a, Choi et al.,2016; Wang et al., 2017; Zhang et al., 2016) and that Gα_o_ can physically interact with mammalian ORs (Scholz et al., 2016b), we looked for functional evidence to implicate Gα_o_ in PI3K-dependent inhibitory signal transduction in mature mouse ORNs. Mice carrying a global deletion of the *Gnao* gene display a variety of defects that include behavioral issues and motor control deficiencies likely resulting from impaired neurogenesis and olfactory system development (Jiang et al., 1998; Choi et al., 2016). Therefore, to test the impact of deletion of *Gnao1* on olfactory signal transduction without possible confounds resulting from widespread issues with olfactory system development, we used a conditional Cre-based knockout (KO) model in which inactivation of *Gnao1* through deletion of exons 5 and 6 is restricted to OMP-positive cells, referred to as c*Gnαo1^-/-^* mice (Chamero et al., 2011; Oboti et al., 2014). Given that OMP expression is restricted to mature ORNs, the OE should develop normally and the impact of Gαo deletion on signaling should be restricted to these cells.

We asked whether depletion of the Gα_o_ protein altered the odor-evoked activity of ORNs by monitoring the responses of acutely dissociated ORNs from C57BI6J and c*Gnαo1^-/-^* mice to a complex odor mixture (H100) in the presence and absence of the PI3K blocker LY294002 (10 μM) (Fig 1A, 1^st^ and 2^nd^ columns, showing type results for 24 ORNs). We predicted that the response evoked by H100 will reflect excitation evoked by one or more components of the mixture that is tempered by inhibition evoked by one or more other components, and that pharmacologically blocking PI3K will result in an increase in the net response magnitude in instances where PI3K-based inhibitory signaling occurs. All cells were also tested with a higher concentration of H100 than the test concentration (Fig 1A, 3^rd^ column). Only those cells showing a 10% or greater response to the higher concentration of H100, indicating their response was not saturated at the test concentration, were subsequently analyzed. The responsiveness of all the ORNs to an IBMX/ forskolin mixture (Fig 1A, 4^th^ column) confirmed the functional integrity of the isolated ORNs.

**Figure 1.**
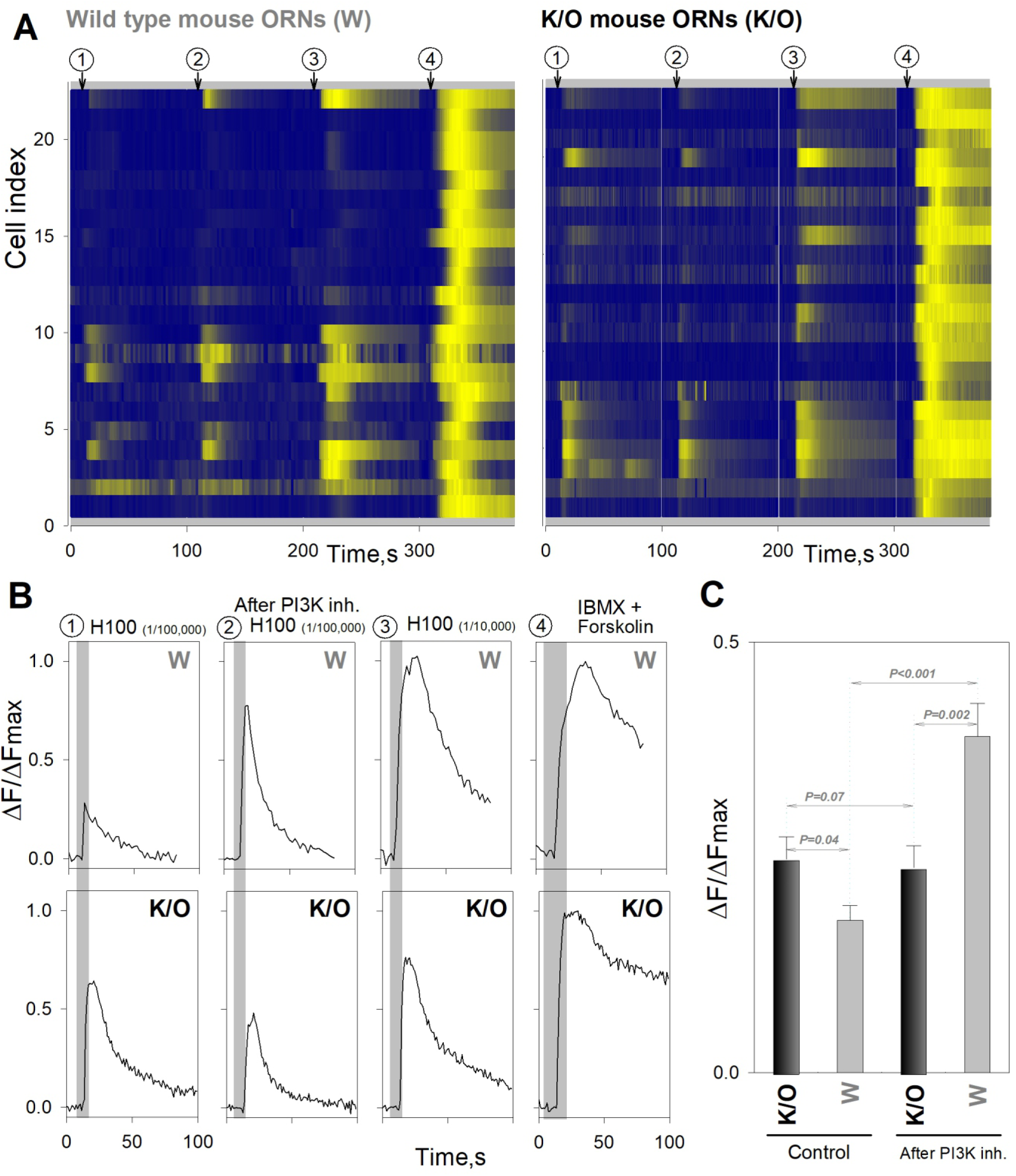
Gα_o_ is required for PI3K-dependent inhibitory signal transduction in mouse ORNs. A. Representative odor-evoked calcium responses of 23 ORNs acutely dissociated from the main olfactory organ of wild type (W) (left panel) and 23 ORNs from knockout (K/O, *cGnαo1^-/-^*) (right panel) mice, before (column 1) and after (column 2) incubation with the PI3K inhibitor LY294002 (LY). Stimulus: Henkel 100 (H100, 1/100,000 dilution). Column 3, responses to a tenfold higher concentration of H100 (1/10,000). Column 4, responses to a mixture of IBMX (50µM) and Forskolin (50µM). The fluorescence intensity traces were normalized to the maximum fluorescent intensity generated in response to IBMX/Forskolin, and then color coded from blue (minimum fluorescent intensity to yellow (maximum fluorescent intensity). Stimulus pulse duration was 5s for columns 1, 2, and 3, and 10s for column 4. B. Representative time/intensity plots of the calcium responses of W type ORN # 21 in A (top row) and K/O type ORN # 6 in A (bottom row). C. Bar graph comparing the average amplitude of the odor-evoked calcium responses of 79 ORNs from K/O mice and 78 ORNs from W mice before and after incubation with LY. Statistical comparisons based on the Mann-Whitney Rank Sum Test.

Our data confirmed the results of previous studies suggesting that ORNs of c*Gnαo1^-/-^* mice maintain their odor responsiveness (Chamero et al., 2011; Oboti et al., 2014). H100 (1:100,000 dilution) evoked a mean response amplitude from ORNs isolated from c*Gnαo1^-/-^* mice of 0.25 ± 0.03 (n = 79 ORNs from 4 mice) (Fig. 1C, 1^st^ bar). The mean response amplitude of the ORNs from c*Gnαo1^-/-^* mice was significantly larger than that observed in the ORNs from WT mice (0.18 ± 0.02, n = 78 ORNs from 5 mice) (Fig. 1C, 2^nd^ bar) (P = 0.04, 1^st^ bar vs 2^nd^ bar), consistent with the inhibitory PI3K signaling pathway not being activated in ORNs from c*Gnαo1^-/-^* mice.

On incubating the ORNs from both the c*Gnαo1^-/-^* and B6 mice with the PI3K blocker LY294002 (10 μM) prior to treatment with H100, no enhancement of the response was observed in the ORNs from c*Gnαo1^-/-^* mice, evoking a normalized mean response amplitude of 0.24 ± 0.03 (n = 79 ORNs from 4 mice) (Fig 1C, 3^rd^ bar) (P = 0.07, 3^rd^ bar vs 1^st^ bar). The lack of change from baseline recordings would be consistent with the inhibitory PI3K signaling pathway not being activated in ORNs from c*Gnαo1^-/-^* mice. In contrast, the response of 39 of 78 (50%) ORNs from WT mice was significantly enhanced by PI3K blockade, evoking a normalized mean response amplitude of 0.39 ± 0.04 (n = 78 ORNs from 5 mice) (Fig 1C, 4^th^ bar) (P = 0.<001, 4^th^ bar vs 2^nd^ bar). The significantly smaller mean response of the ORNs from c*Gnαo1^-/-^*mice (P = 0.002, 3^rd^ bar vs 4^th^ bar) than that observed in the ORNs from WT mice would be consistent with the inhibitory PI3K signaling pathway not being activated in ORNs from c*Gnαo1^-/-^* mice. Presumably odor stimulation should not have activated PI3K signaling in either group of cells treated with LY294002. Thus, finding that blockade of PI3K in WT ORNs resulted in a normalized mean response that was significantly larger than that of ORNs from the c*Gnαo1^-/-^* mice (Fig. 1C, 4^th^ bar vs 3^rd^ bar) could potentially indicate Gα_o_-independent activation of PI3K. However, that is not likely since blockade of PI3K had no effect on the response of ORNs from the c*Gnαo1^-/-^*mice (Fig. 1C, 3^rd^ vs 1^st^ bars), suggesting the magnitude of the response in WT ORNs post-blockade reflects the dynamics of action of the drug when PI3K is activated. Collectively, these findings are consistent with the hypothesis that Gα_o_ is functionally upstream of PI3K in the context of inhibitory transduction in mature ORNs.

### Gα_o_ expression in the OE in mice carrying an OMP-Cre based deletion of Gnao1

The loss of PI3K-based inhibition of excitatory signaling in odorant sensitive ORNs in the c*Gnαo1^-/-^* mice implies that Gα_o_ localizes to the olfactory cilia where transduction occurs and that it is reduced in the c*Gnαo1^-/-^* mice. A previous study validated reduced Gnao1 gene expression in the c*Gnαo1^-/-^*mice, but did not find reduced Gnao1 gene expression the total OE (Chamero et al., 2011). Here we show deletion of exons 5 and 6 occurs in the OE at the genomic level using PCR with primers spanning this region (Fig 2A; Choi et al., 2016). The recombined *Gnao1* gene is present as a smaller fragment amplified from DNA isolated from the OE and VNO, but is at low to undetectable levels in the olfactory bulb where there is no OMP expression. Recombination is absent in B6 mice. *In situ* hybridization targets *Gnao1* expression to the mature ORN (OMP-expressing) layer of the OE (Fig 2B; Heron et al., 2013; Saraiva et al., 2015; Choi et al., 2016). Since *Gnao1* mRNA is abundant in ORNs (Heron et al., 2013; Omura and Mombaerts, 2014; Saraiva et al., 2015; Wang et al., 2017), at least some of the recombination potentially could be ascribed to mature ORNs. Immuno-labeling cryosections of the OE localized expression of the Gα_o_ protein to the axon bundles mature (OMP-expressing) ORNs but was unable to localize expression of the Gα_o_ protein to the distal compartments and/or cilia (data not shown).

**Figure 2.**
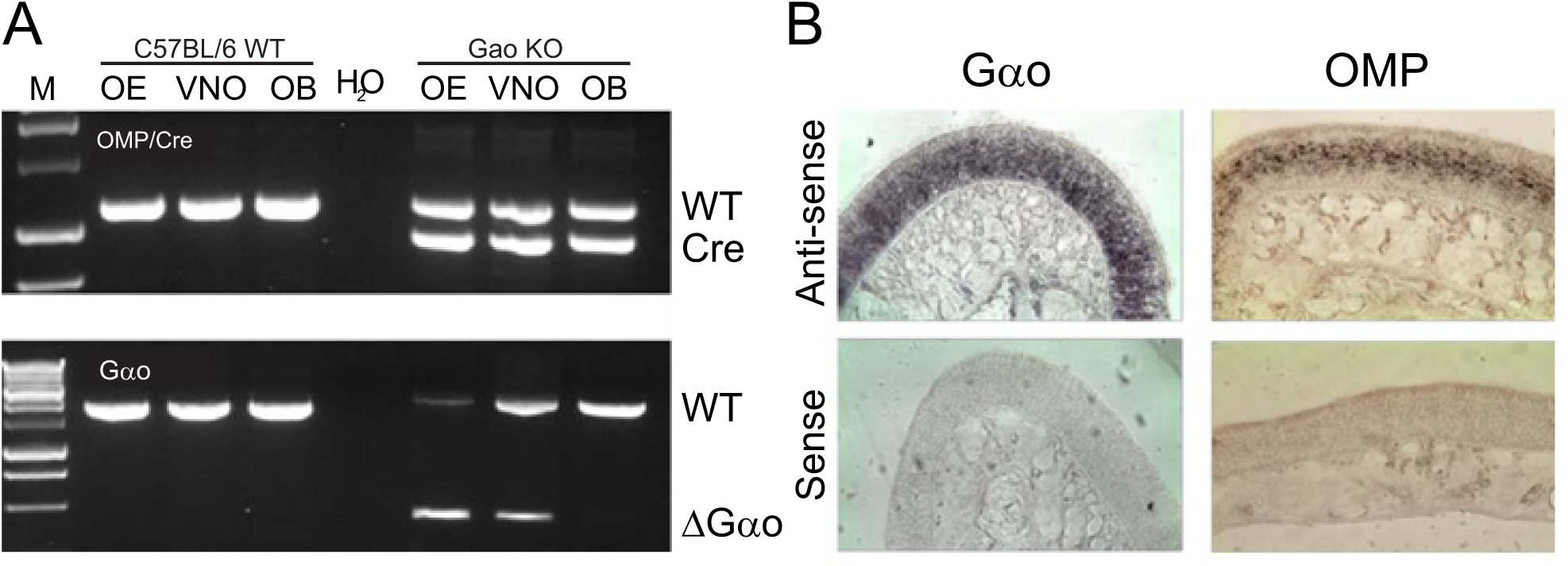
OMP-Cre mediated Gnao1 deletion in the olfactory epithelia of *cGnαo1^-/-^* knockout mice. A. Genomic DNA extracted from the OE, VNO, and OB was examined by PCR for recombination of floxed alleles. C57BL/6 WT (WT) mice were used for comparison. In the KO mice, both the WT and OMP-Cre (Cre) alleles are detected. The recombined Gnao1 (ΔGαo) allele is detected in the OE and VNO of the KO mice as indicated by the smaller fragment in the lower panel. B. Comparison of Gnao1 (Gαo) and OMP expression in the OE of B6 mice by *in situ* hybridization of cryosection from B6 mice. OMP expression is restricted to the mature ORNs.

We instead determined whether ectopically expressed Gα_o_ can be trafficked to the cilia (McEwen et al., 2008; McIntyre et al., 2015). Mice were intra-nasally injected with adeno-associated virus carrying fluorescently tagged Gα_o_ (Hynes et al., 2004) (AAV Gα_o_:mcherry). *En face* imaging revealed Gα_o_:mCherry in the cilia of transduced ORNs (Fig 3A). In ORNs transduced with adenovirus carrying Gα_o_:GFP, GFP expression co-localized with the ciliary protein Arl13b (Fig. 3C), indicating that these results are not dependent on the identity of the fluorescent tag inserted into Gα_o_ or on the viral vector used for infection. Given that the OE is composed of multiple types of chemosensory cells (e.g., Munger, 2009), we then asked whether Gα_o_ could localize to the cilia of ORNs expressing ORs known to couple to Gα_olf_. Using mice expressing tauGFP under the control of the SR1 OR gene, we found AAV expressed Gα_o_:mCherry localized GFP^+^ to the cila (Fig. 3B), suggesting it is not excluded from canonical ORNs

**Figure 3.**
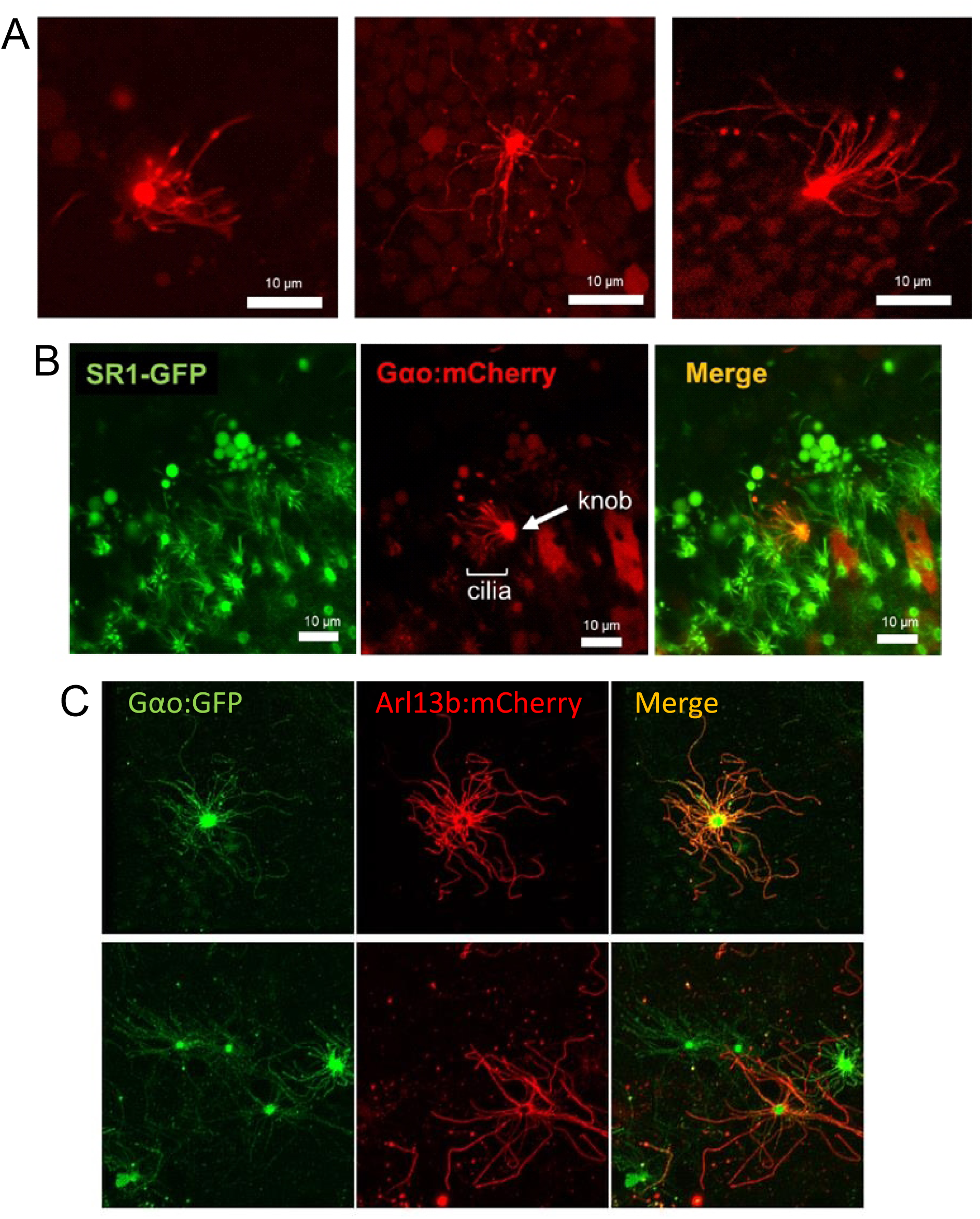
Ectopically expressed Gα_o_ can enter the cilia of mammalian ORNs. *En face* imaging of ORNs expressing Gα_o_ internally tagged with mCherry or GFP. A. Three examples of AAV infected ORNs ectopically expressing Gα_o_:mCherry. B. Gα_o_:mCherry can be co-localized to SR1-GFP+ ORNs. mCherry expression is found throughout infected ORNs including in the dendritic knobs and cilia. Scale bars represent 10 μM. C. AV infected ORNs of C57BL/6 mice ectopically expressing Gα_o_:GFP and Arl13b:mCherry. Gα_o_ expression overlaps with that of Arl13b indicating that ciliary localization of the ectopically expressed protein does not depend on the vector or tag.

### Gα_o_ enhances odorant-evoked coupling of a mammalian OR isolated from native ORNs responsive to an identified opponent odorant pair in HEK293T cells

PI3K dependent inhibitory signaling has been demonstrated in both rats and mice (e.g., Brunert et al., 2010; Ukhanov et al., 2010). Several opponent (excitatory/inhibitory) odorant pairs have been identified for rat ORNs (e.g., Ukhanov et al., 2010; 2011), and here use one of those pairs to assess whether a single mammalian OR can activate both PI3K signaling through Gα_o_ and ACIII signaling through Gα_olf_.

We first measured the calcium signal in acutely dissociated rat ORNs evoked by octanol (OOL, 50 μM) both alone and in combination with citral (CIT, 100 μM). In a subset of OOL-responsive cells, co-application of CIT reduced the peak Ca^2+^ response by 5-fold on average (Fig, 4A). Pre-incubation of the cells with the PI3Kβ and -γ isoform specific blockers TGX221 and AS252424 (200 nM each) rescued the Ca^2+^ response (Fig, 4A), indicating that the antagonism was not the result of direct competition of the odorants for the binding site, but rather activation of the opponent inhibitory PI3K signaling pathway. Individual ORNs with this response profile were collected (Fig, 4B) for single cell RT-PCR using degenerate primers based on conserved regions of mammalian OR sequences (Touhara et al., 1999). Prior to OR amplification, the samples were tested for olfactory marker protein (OMP) expression to ensure that they were mature ORNs (Barber et al., 2000) and with β-actin primers to avoid testing those with detectable genomic DNA contamination (Chan et al., 1997). From a total of ten functionally delimited rat ORNs that met these requirements, we recovered three rat ORs (Olr1845, two ORNs; Olr1479, two ORNs; Olr1231, one ORN; no OR amplified, five ORNs) and cloned the full length sequences for heterologous expression under the control of a CMV promoter with Lucy and Rho tags to enhance their surface expression (Shepard et al., 2013).

**Figure 4.**
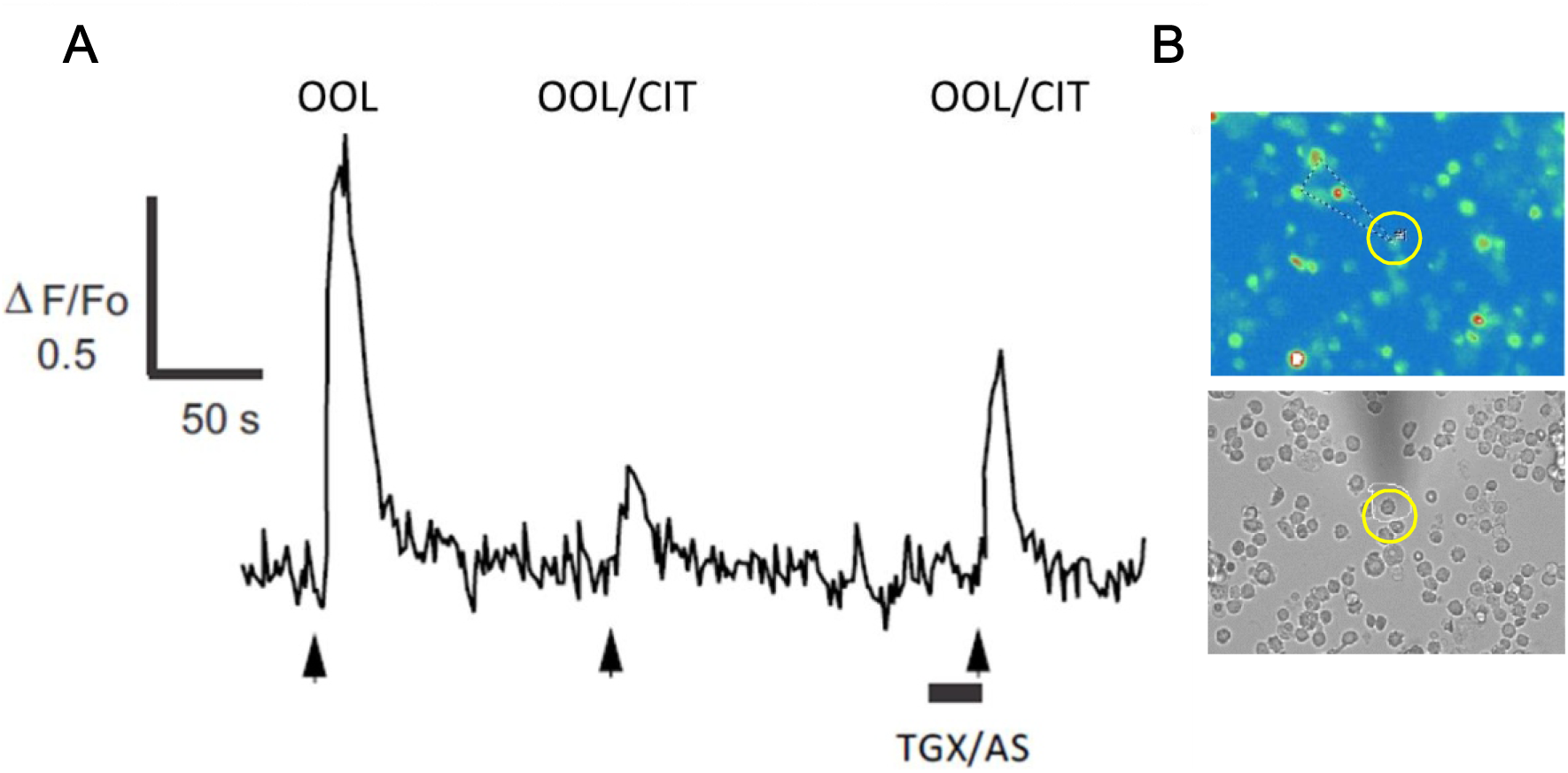
Identification of ORNs responsive to the antagonistic odorant pair octanol and citral for single cell RT-PCR. A. Fluo-3 calcium imaging of dissociated rat ORNs was used to identify single cells for RT-PCR. Representative recording of the somatic Ca2+ response from one of ten rat ORNs activated by octanol (OOL; 50 μM) in which citral (CIT; 100 μM) inhibited the response and pretreatment with PI3K inhibitors TGX221 and AS252424 (TGX/AS; 200 nM each) partially relieved the antagonism. B. Image of an ORN identified by calcium imaging prior to collection for single cell RT-PCR.

We then tested the function of the receptors in a pCRE-SEAP assay by co-expressing them with Gα_olf_, RTP1s and Ric8b in HEK293T cells along with a cAMP reporter gene (Durocher et al., 2000; Zhuang and Matsunami, 2007). The cAMP reporter plasmid pCRE-SEAP expresses secreted alkaline phosphatase (SEAP) in response to cAMP binding to cAMP response elements (CRE). Odorants were added at 48 hours post transfection and SEAP activity was measured 20 hours later. All results represent at least three independent replicate experiments. We focused on Olr1845, which responded consistently to OOL in a dose-dependent manner (Fig 5A). The other receptors did not respond consistently and will require further optimization to determine whether they show similar ligand profiles. Olr1845 did not produce measurable responses to other single odorants tested (250 μM) including vanillin, eugenol, and isovalaric acid (data not shown). Olr1845 did not respond to 75 μM CIT alone, but 75 μM CIT suppressed the response to OOL in a graded manner (Fig 5B). Control experiments in which cells were transfected with all of the signaling co-factors, except, Olr1845, and tested in parallel did not show changes in SEAP activity when stimulated with OOL alone or in combination with CIT (Fig. 5B, inset). We then tested the mouse OR OR261-1 (Olfr447), known to respond to OOL (Saito et al., 2009), and confirmed its response to OOL in a dose-dependent manner (Fig 5C). In contrast to Olr1845, OR261-1 responded to 75 μM CIT alone (Fig 5D), which was not affected by increasing concentrations of OOL. This result indicates that not all receptors that respond to OOL respond to CIT, or a mix of CIT and OOL, in the same manner. The experimental results with Olr1845 also serve as a positive control, allowing us to assign the effects seen with Olr1845 to that receptor and not one inherent in the heterologous cell.

**Figure 5.**
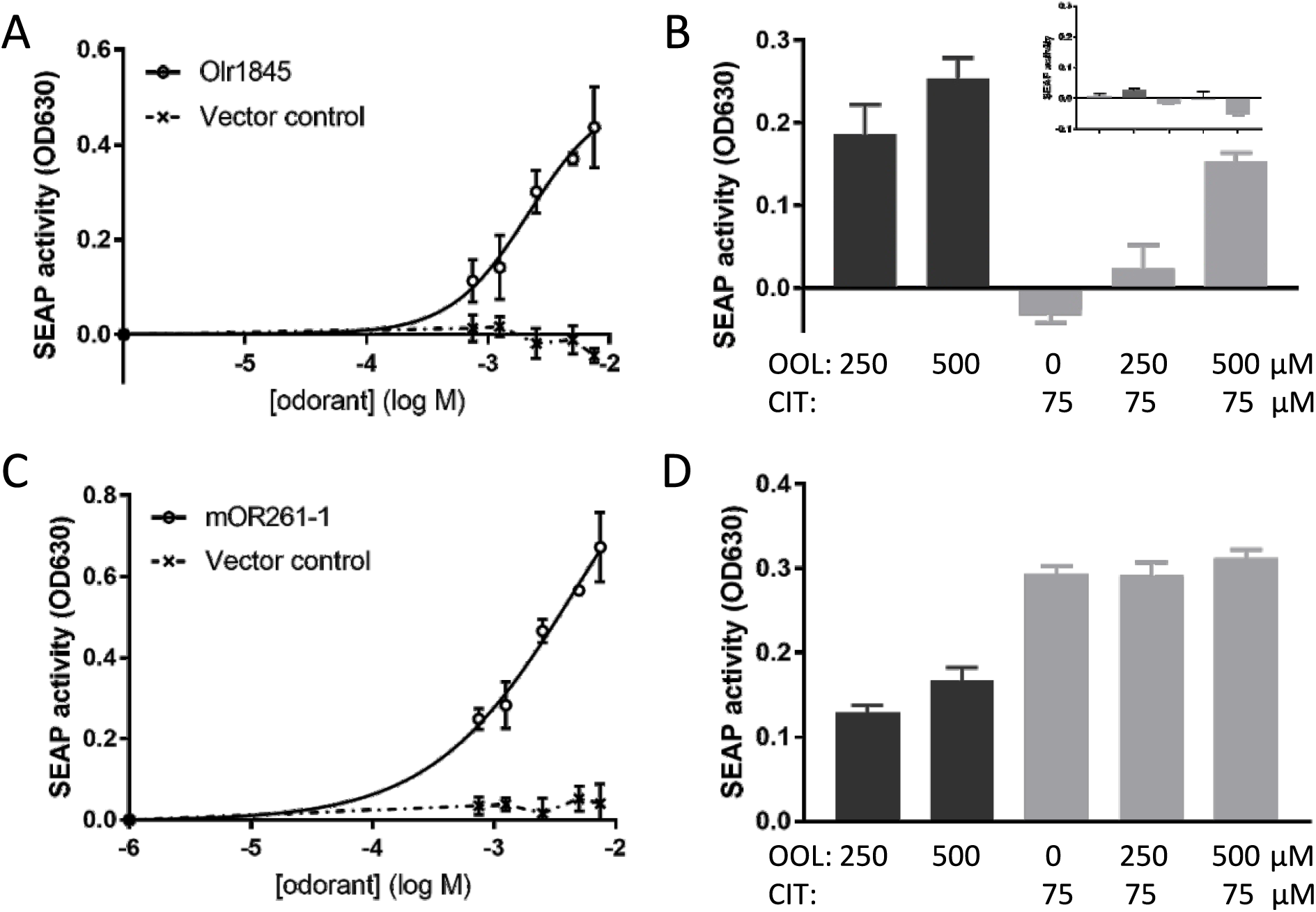
Gαolf and ACIII activation by rat Olr1845 in response to octanol and citral. A. Line graph showing that rat Olr1845 responds in a dose dependent manner to OOL in a pCRE-SEAP assay. Response to OOL is denoted by open circles. B. Bar graph showing that the cAMP response of rat Olr1845 to OOL at the concentrations indicated (dark bars) was reduced in a graded manner when the odorant was presented in a binary mixture with CIT at the concentration indicated (light bars). Inset: Bar graph showing the response of cells not expressing an OR tested in the same experimental paradigm. C. Line graph showing that a different mouse OR (mOR261-1) also responds in a dose dependent manner to OOL in a pCRE-SEAP assay (open circles). D. Bar graph showing that in contrast to B, the cAMP response of mouse OR261-1 to OOL at the concentrations indicated. (dark bars) was actually enhanced, i.e., shows additivity, when the odorant was presented in binary mixture with CIT at the concentration indicated (light bars). Data are presented as SEAP activity (OD630) -/+ SEM representing at least three independent replicate experiments. Response of cells to DMSO has been subtracted in all cases.

To determine whether Gα_o_ enhances the odorant-evoked coupling of Olr1845 to PI3K, we used an ELISA specific for PIP3, the primary product of PI3K activation *in vivo* (Ukhanov et al., 2010). We first co-expressed Olr1845 with RTP1s in HEK293T cells, relying on the endogenous G proteins and associated chaperones. At 48 hours post-transfection, a 30 sec treatment of the cells with CIT or OOL (500 μM), increased the level of PIP3 by 49.52 ± 3.71 pmol and 25.78 ± 7.02 pmol, respectively, (n = at least 3 independent replicates) above that in response to carrier (DMSO) treatment alone. This indicates that Orl1845 can activate the PI3K pathway in the heterologous system and that CIT is a stronger PIP3-dependent agonist. The response to CIT is significantly higher than that to OOL (Fig. 6, 1^st^ pair of bars, P=0.04).

**Figure 6.**
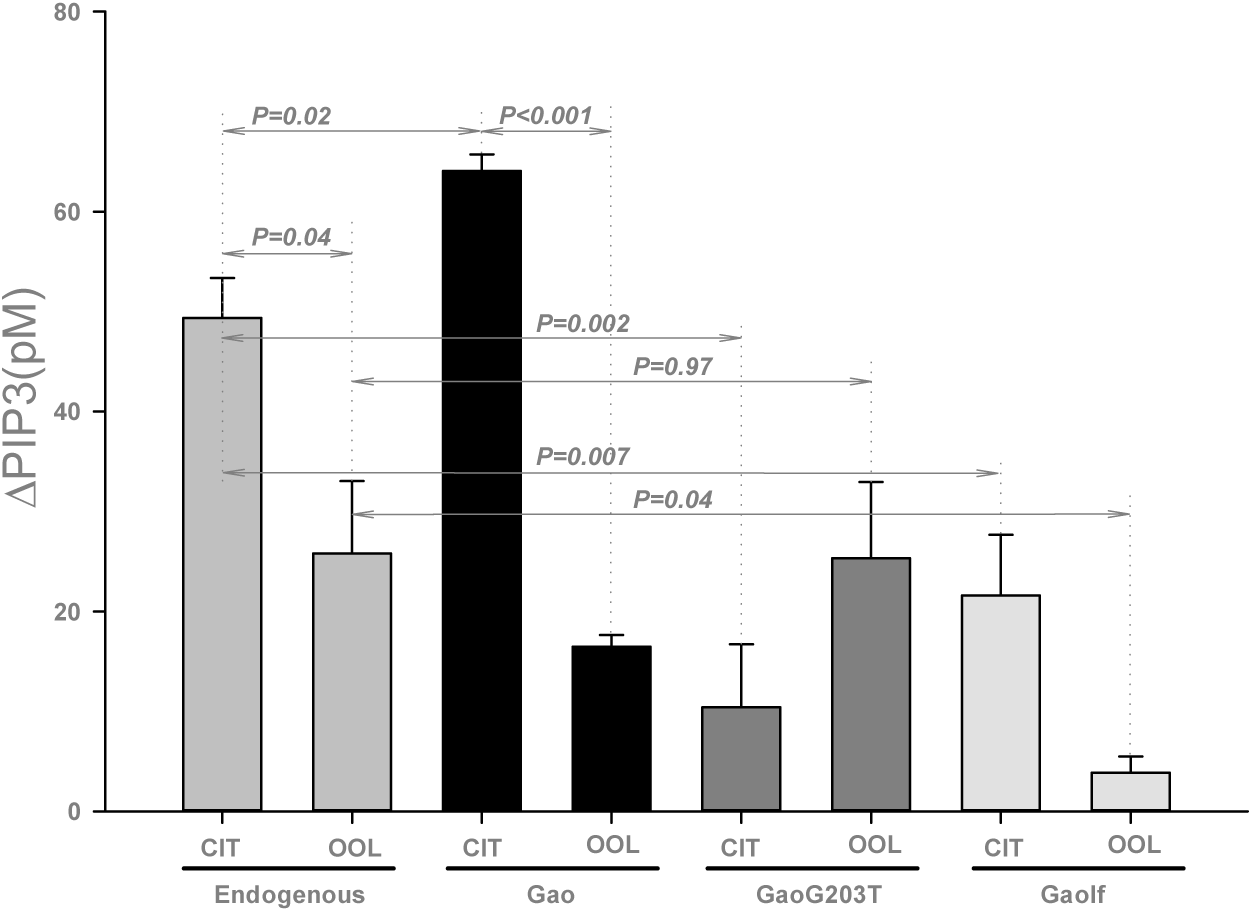
PI3K activation by rat Olr1845 in response to octanol and citral. Bar graph showing the elevation of endogeneous PIP3 in HEK293T cells transfected with rat Olr1845 in response to citral (CIT; 500 μM) and octanol (OOL; 500 μM). PI3K activity was measured by a PIP3 ELISA at 48 hours post-transfection in response to a 30 sec odorant exposure. The PIP3 level in DMSO-treated control cells is subtracted in all instances. The receptor was either expressed alone (endogeneous, 1^st^ pair of bars), with Gα_o_ (2^nd^ pair of bars), with Gα_oG203T_ (3^rd^ pair of bars), or with Gα_olf_ (4^th^ pair of bars) Data are presented as change in pmol PIP3 ± SEM, representing at least three independent replicates. Probabilities for the various comparisons listed in the text are indicated by the horizontal lines. Statistical comparison based on the Student’s t test.

We then independently co-expressed three different Gα subunits together with Olr1845, each with at least three independent replicates. Gα_o_ overexpression significantly enhanced the increase in PI3K activation in response to CIT, resulting in a PIP3 level of 64.16 ± 1.44 pmol (Fig. 6, 1^st^ bars in the 1^st^ and 2^nd^ pair of bars, P= 0.02). This suggests that Gα_o_ plays a role in mediating PI3K activation in the heterologous system. Again, the response to CIT was significantly higher than that to OOL (Fig. 6, 2^nd^ pair of bars, P=0.001). To test whether the increase in PI3K signaling resulted from the functional activity of Gα_o_, we co-expressed a Gα_o_ gene carrying a G203T mutation that is predicted to decrease the ability of the protein to turn over GDP and GTP (Slepak et al., 1993) and attenuate its ability to activate downstream signaling. Unlike native Gα_o_, the mutated G protein subunit resulted in a lower level of PI3K activation by CIT (10.54 ± 3.46 pmol) in comparison to cells expressing only endogenous G proteins, actually significantly decreasing it, and no change in the level of PI3K activation by OOL (25.29 ± 7.62 pmol), indicating that the activity of Gα_o_ is required for this process (Fig. 6, 3^rd^ pair of bars vs 1^st^ pair of bars, P = 0.002; 0.97, respectively). Gα_olf_ co-expression, which enhances the cAMP response of ORs (Zhuang and Matsunami, 2007), also resulted in lower levels of PI3K activation by both CIT and OOL (21.60 ± 4.17 pmol and 3.75 ± 1.24 pmol, respectively) in comparison to cells expressing only endogenous G proteins, actually significantly decreasing them (Fig. 6, 4^th^ pair of bars vs 1^st^ pair of bars, P = 0.007; 0.04, respectively). This latter effect potentially reflects sequestration of the necessary Gβγ subunits from endogenous G proteins (Hippe et al., 2013).

## Discussion

Our findings potentially associate a second G protein, Gα_o_, with inhibitory signalling in mammalian olfactory transduction by first showing that odor evoked phosphoinositide 3-kinase (PI3K)-dependent inhibition of signal transduction is absent in the native ORNs of mice carrying a conditional OMP-Cre based knockout of Gα_o_. Finding that conditional Gα_o_ deletion eliminates odor evoked, PI3K-dependent inhibition in dissociated ORNs argues that Gα_o_ is functionally upstream of PI3K. We then identify an OR from native rat ORNs that are activated by octanol through cyclic nucleotide signaling and inhibited by citral in a PI3K-dependent manner. We show that the OR activates cyclic nucleotide signaling and PI3K signaling in a manner that reflects its functionality in native ORNs. Collectively these findings raise the interesting possibility that a mammalian OR can interact with two different G proteins in a functionally identified, ligand-dependent manner.

We argue that functional data obtained in acutely dissociated ORNs implicate Gα_o_ in mediating inhibitory olfactory signal transduction. Acutely dissociating ORNs destroys the normal polarity of the cells, exposing the entire cell to odors and allowing that Gα_o_-dependent activation of PI3K is not related to transduction per se. Two findings counter this possibility. First, odorant sensitivity is predominately, if not entirely, localized to the cilia/dendritic knob in dissociated vertebrate (salamander) ORNs (Lowe and Gold, 1991) and the dissociated mammalian ORNs typically retain at least part of their ciliary complement in our hands. Second, the pharmacological effect of blocking PI3K-dependent inhibition seen here in the dissociated ORNs occurs with the same dynamics in ORNs in the intact OE where the normal polarity of the cells targets signaling to the cilia/dendritic knob, and where patch-clamping dendritic knobs shows that PI3K-dependent inhibition acts with msec resolution, setting the peak frequency and the latency of the train of action potentials evoked by an odor mixture (Ukhanov et al., 2010). Thus, we have no reason to assume the functional data are biased by using acutely dissociated ORNs. However, given we did not use littermate controls the possibility remains that the functional data reflect strain differences, which will require further experimentation to resolve.

Implicating Gα_o_ in the activation of PI3K is consistent with evidence that PI3K signaling is sensitive to pertussis toxin, which is indicative of Gα_i/o_-dependency in other systems (e.g. Orr et al., 2002; Banquet et al., 2011; Hadi et al., 2013), as well as with evidence that Gα_o_ can signal through interactions of its associated βγ subunits with downstream effectors (Wettschureck, 2005; Steiner et al., 2006; Bondar and Lazar, 2014). Published evidence shows that class 1B PI3Kγ is expressed in mouse ORNs, that PI3Kγ-deficient mice show almost a complete lack of odorant-induced PI3K activity in their OE, and that the ORNs of PI3Kγ-deficient mice show reduced sensitivity to PI3K mediated inhibition (Brunert et al., 2010). The γ catalytic subunit is thought to exclusively associate with the regulatory subunits that mediate binding to the Gβγ subunit of heterotrimeric G proteins (e.g., Rameh and Cantley, 1999), which in our case would be Gα_o_ activated by the OR.

Gα_o_ also mediates PLC signaling in mature ORNs (Schandar et al., 1998), as well as extracellular-signal-regulated kinase (ERK) signaling by a heterologously expressed OR (Bush et al., 2007) that is associated with cell survival and apoptosis. Since both the PLC and ERK pathways have been associated with PI3K-dependent signaling networks in other systems, temporally distinct waves of Gα_o_ activated PI3K (e.g., Jones et al., 1999; Goncharova et al., 2002) in ORNs could potentially mediate transduction as well as slower activation of cell survival and/or apoptotic pathways. There has been a published report for a Gα_o_-mediated alternate, cyclic nucleotide-independent, PI3K-independent signaling pathway in mammalian ORNs that targets a downstream Cl^-^ conductance and presumably leads to an excitatory efflux of Cl^-^ (Scholz et al., 2016b). The functional significance of this pathway is unclear, but as the authors suggest, this pathway may be developmentally important since it appears to be associated with immature ORNs. Further work will be required to relate this finding to the Gα_o_-mediated inhibitory signaling in mature ORNs proposed herein.

We show that excitation and PI3K-dependent inhibition can be mediated by the same OR when expressed heterologously, and that the antagonism is not the result of direct competition for a common binding site. As in the native ORNs from which the rat OR Olr1845 was cloned, OOL acts as an excitatory ligand and CIT as an inhibitory ligand for Olr1845 in a cAMP assay when heterologously expressed. This does not occur when the cells were transfected with the co-factors in the absence of the OR, nor when the cells were transfected with a different OR responsive to the same two ligands. We also show that CIT is a stronger activator of PI3K than is OOL in a PIP3 assay when Olr1854 is heterologously expressed. Heterologous readouts of OR activation are slow in comparison to transduction, but similar assays reflect the ligand specificity of other ORs tested *in vivo* (Tsuboi et al., 2011), supporting our argument that Olr1845 appears to be capable of directing the pattern of activation elicited by an opponent pair of ligands through two different signaling pathways. This finding for a mammalian OR is consistent with the ability of single insect olfactory receptors to similarly direct the pattern of activation of an ORN in studies using the ‘empty neuron’ approach (Hallem et al., 2004). Whether all ligands in the molecular receptive range of a given OR can activate PI3K, only to different extents, with the stronger PI3K-dependent agonists being the effective inhibitory ligands for the OR in question, as potentially suggested by Fig. 6, remains for future research. We focused on Olr1845, which allows that our finding could be idiosyncratic for Olr1845 or the OOL/CIT odorant pair, but evidence that Gα_o_ can interact with other mouse ORs (Scholz et al., 2016b), as well as evidence that PI3K-dependent inhibition can be activated by a wide range of conventional odorants in native rat ORNs (Ukhanov et al., 2013), including other opponent odorant pairs (Ukhanov et al., 2011), argues for the generality of this finding across at least a subset of ORs.

Assuming the OR and Gα_o_ interact, the assumption would be they interact in the transduction (ciliary) compartment. As noted, both immature and mature ORNs express Gα_o_ (Mayer et al., 2009; Keydar et al., 2013; Heron et al., 2013; Nickell et al., 2012; Omura and Mombaerts, 2014; Saraiva et al., 2015; Scholz et al., 2016a, Choi et al.,2016; Wang et al., 2017; Zhang et al., 2016). It remains to be determined, however, whether the protein routinely localizes to the ciliary compartment, notwithstanding limited evidence for positive IHC staining for Gα_o_ in the distal compartments of OMP^+^ ORNs in *Gnao1*^+/+^ mice (Choi et al., 2016) and *Olfr73*-positive ORNs (Scholz et al., 2016a). The fact that we could show there appears to be no barrier excluding Gα_o_ from the ciliary compartment is consistent with ciliary expression since cilia are known to largely exclude non-resident proteins (McEwen et al., 2008; McIntyre et al., 2015), although this demonstration leaves open the question of constitutive expression of Gα_o_ in the ciliary compartment. While Gα_o_ is expressed in sustentacular cells (SUSs) (unpublished observations), the possibility that it interacts with signaling in ORNs via ephaptic coupling (Su, et al., 2012) is not consistent with our physiological results obtained in acutely dissociated ORNs. Nor is it consistent with the absence of any evidence that mammalian ORNs and SUSs are grouped in stereotyped functional combinations that would be required to explain the observed ligand specificity.

GPCRs are increasingly appreciated to sequentially activate multiple G proteins such that the outcome of activation does not depend solely on the receptor identity but rather is influenced by extracellular factors such as the range of ligands present, as well as by intracellular factors including the abundance and localization of the G proteins present (e.g., Mashuo et al., 2015 and reviewed in Lohse and Hofmann, 2015; Latorraca et al., 2016). Such ‘functional selectivity’ (e.g., Luttrell, 2014; Smrcka, 2015) is a key characteristic of allosteric modulation in GPCRs (Christopoulis, 2014). Given that OR-ligand interaction is thought to be ‘fast and loose’ (Bhandawat, 2005) and growing evidence for loose allosteric coupling of the agonist binding site and the G protein coupling interface in GPCRs (e.g., Lohse and Hofmann, 2015; Manglik et al., 2015; Wingler et al., 2019), a given OR could interact with both G protein isoforms without implying concurrent activation by a given odorant or the need for simultaneous coupling of the OR to both Gα_olf_ and Gα_o_. Brief activation of the OR by a PI3K-dependent inhibitory ligand, for instance, could release pre-bound Gα_olf_ while resulting in a more favorable structure for binding to Gα_o_. The fact that not all G proteins work *in vivo* by having the heterotrimers physically dissociate (e.g., Digby et al., 2006) could provide specificity for signals mediated by the βγ dimer, as in the present context, and avoid confound in the origin of the βγ dimer. However, the idea that ligand-bound GPCRs interact with and activate G proteins (e.g., Audet et al., 2012) is being replaced by emerging evidence that the GPCR and G protein are preassembled into protein complexes in which the G protein influences ligand affinity (e.g., DeVree et al., 2016; Venkatakrishnan et al., 2016). Thus, a subset of the OR expressed by a given cell could be primed for activation of Gα_o_, and in turn PI3K inhibitory signaling. Understanding how functional selectivity in ORs could play out at the molecular level awaits further understanding of GPCR signaling in general.

In summary, our findings lay the groundwork to explore the interesting possibility that ORs can interact with two different G proteins in a functionally identified, ligand-dependent manner to mediate opponent signaling in mature mammalian ORNs. Going forward, the primary challenge will be to understand the expression pattern of Gα_o_, and potentially other G proteins, in addition to Gα_olf_ in the transduction compartment. If Gα_o_ is present in cilia, but perhaps at lower levels or not in all neurons, this may be better approached through genetic methods to label and visualize the endogenous protein. The possibility that ORs can interact with multiple G proteins in a functionally identified, ligand-dependent manner in the context of transduction would be a paradigm-shift in our understanding of how the olfactory periphery sets the combinatorial pattern considered to be the basis of odor recognition and discrimination.

## Acknowledgements

We thank Drs. Frank Zufall and Trese Leinders-Zufall (Saarland U., Homburg, Germany) for kindly providing the breeding stock for the c*Gnαo1^-/-^* mice, Dr. Sergei Zolotukhin (U. Florida) for assistance and guidance with adeno-associated virus infection, Dr. Hanns Hatt (Ruhr U., Bochum, Germany) for kindly providing the odorant mixture Henkel 100, and Leanne Adams (U. Florida) for technical assistance.

## Author Contributions

EAC, KU, YUB, JCM, and JRM designed the experiments. EAC, KU, YUB carried out the experiments and analyzed the data. EAC, BWA, and JCM prepared the manuscript. All authors discussed and contributed to writing the manuscript.

## References

Audet M et al. (2012) Restructuring G protein-coupled receptor activation. Cell 151:14–23.

Balla T, Szentpetery Z, Kim YJ (2009) Phosphoinositide signaling: new tools and insights. Physiology (Bethesda) 24:231–244.

Banquet S, Delannoy E, Agouni A, Dessy C, Lacomme S, Hubert F, Richard V, Muller B, Leblais V (2011) Role of Gi/o-Src kinase-PI3K/Akt pathway and caveolin-1 in β2-adrenoceptor coupling to endothelial NO synthase in mouse pulmonary artery. Cell Signal 23:1136–1143.

Barber RD, Jaworsky DE, Yau KW, Ronnett G V (2000) Isolation and in vitro differentiation of conditionally immortalized murine olfactory receptor neurons. J Neurosci 20:3695–3704.

Bell GA, Laing DG, Pahhuber H (1987) Odor mixture suppression: evidence for a peripheral mechanism in human and rat. Brain Res. 426:8–18.

Belluscio L, Gold GH, Nemes A, Axel R (1998) Mice deficient in Golf are anosmic. Neuron 20:69–8.

Bhandawat V (2005) Elementary response of olfactory receptor neurons to odorants. Science (80-) 308:1931–1934.

Bondar A, Lazar J (2014) Dissociated G GTP and G Protein subunits are the major activated form of heterotrimeric Gi/o proteins. J Biol Chem 289:1271–1281.

Brady JD, Rich ED, Martens JR, Karpen JW, Varnum MD, Brown RL (2006) Interplay between PIP3 and calmodulin regulation of olfactory cyclic nucleotide-gated channels. Proc Natl Acad Sci U S A 103:15635–15640.

Brunert D, Klasen K, Corey EA, Ache BW (2010) PI3Kgamma-dependent signaling in mouse olfactory receptor neurons. Chem Senses 35:301–308.

Buck L, Axel R (1991) A novel multigene family may encode odorant receptors: A molecular basis for odor recognition. Cell 65:175–187.

Bush CF, Jones S V, Lyle AN, Minneman KP, Ressler KJ, Hall RA (2007) Specificity of olfactory receptor interactions with other G protein-coupled receptors. J Biol Chem 282:19042–19051.

Cain WS. (1974) Odor intensity – mixtures and masking. B Psychonomics Soc 4:244–244.

Chamero P, Katsoulidou V, Hendrix P, Bufe B, Roberts R, Matsunami H, Abramowitz J, Birnbaumer L, Zufall F, Leinders-Zufall T (2011) G protein G(alpha)o is essential for vomeronasal function and aggressive behavior in mice. Proc Natl Acad Sci U S A 108:12898–12903.

Chan SL, Perrett CW, Morgan NG (1997) Differential expression of alpha 2-adrenoceptor subtypes in purified rat pancreatic islet A- and B-cells. Cell Signal 9:71–78.

Choi J-M, Kim S-S, Choi C-I, Cha HL, Oh H-H, Ghil S, Lee Y-D, Birnbaumer L, Suh-Kim H (2016) Development of the main olfactory system and main olfactory epithelium-dependent male mating behavior are altered in G_o_-deficient mice. Proc Natl Acad Sci 113:10974–10979.

Christopoulis A (2014) Advances in G protein-coupled receptor allostery: from function to structure. Mol Pharmacol 86:463–78.

DeVree BT, Mahoney JP, Vélez-Ruiz GA, Rasmussen SGF, Kuszak AJ, Edwald E, Fung J-J, Manglik A, Masureel M, Du Y, Matt RA, Pardon E, Steyaert J, Kobilka BK, Sunahara RK (2016) Allosteric coupling from G protein to the agonist-binding pocket in GPCRs. Nature 535:182–186.

Digby GJ, Lober RM, Sethi PR, Lambert NA (2006) Some G protein heteromers physically dissociate in living cells. Proc Natl Acad Sci 103:17789–17794.

Durocher Y, Perret S, Thibaudeau E, Gaumond MH, Kamen A, Stocco R, Abramovitz M (2000) A reporter gene assay for high-throughput screening of G-protein-coupled receptors stably or transiently expressed in HEK293 EBNA cells grown in suspension culture. Anal Biochem 284:316–326.

Firestein, S, Shepherd, G (1992) Neurotransmitter antagonists block some odor responses in olfactory receptor neurons. Neuroreport 3:661–664.

Goncharova EA, Ammit AJ, Irani C, Carroll RG, Eszterhas AJ, Panettieri RA, Krymskaya VP (2002) PI3K is required for proliferation and migration of human pulmonary vascular smooth muscle cells. Am J Physiol Lung Cell Mol Physiol 283:L354–L363.

Hadi T, Barrichon M, Mourtialon P, Wendremaire M, Garrido C, Sagot P, Bardou M, Lirussi F (2013) Biphasic Erk1/2 activation sequentially involving Gs and Gi signaling is required in beta3-adrenergic receptor-induced primary smooth muscle cell proliferation. Biochim Biophys Acta - Mol Cell Res 1833:1041–1051.

Hallem EA, Ho MG, Carlson JR (2004) The molecular basis of odor coding in the *Drosophila* antenna. Cell 117: 965–979.

Heron PM, Stromberg AJ, Breheny P, McClintock TS (2013) Molecular events in the cell types of the olfactory epithelium during adult neurogenesis. Mol Brain 6:49.

Hynes TR, Hughes TE, Berlot CH (2004) Cellular localization of GFP-tagged alpha subunits. Meth Mol Biol 237:233–46.

Ishii T, Omura M, Mombaerts P (2004) Protocols for two- and three-color fluorescent RNA in situ hybridization of the main and accessory olfactory epithelia in mouse. J Neurocytol 33:657–669.

Jiang M, Gold MS, Boulay G, Spicher K, Peyton M, Brabet P, Srinivasan Y, Rudolph U, Ellison G, Birnbaumer L (1998) Multiple neurological abnormalities in mice deficient in the G protein Go. Proc Natl Acad Sci U S A 95:3269–3274.

Jones SM, Klinghoffer R, Prestwich GD, Toker A, Kazlauskas A (1999) PDGF induces an early and a late wave of PI 3-kinase activity, and only the late wave is required for progression through G1. Curr Biol 9:512–521.

Keydar I, Ben-Asher E, Feldmesser E, Nativ N, Oshimoto A, Restrepo D, Matsunami H, Chien M-S, Pinto JM, Gilad Y, Olender T, Lancet D (2013) General olfactory sensitivity database (GOSdb): candidate genes and their genomic variations. Hum Mutat 34:32–41.

Kim SY, Mammen A, Yoo S-J, Cho B, Kim E-K, Park J-I, Moon C, Ronnett G V (2015a) Phosphoinositide and Erk signaling pathways mediate activity-driven rodent olfactory sensory neuronal survival and stress mitigation. J Neurochem 134:486– 498.

Kim SY, Yoo S-J, Ronnett G V, Kim E-K, Moon C (2015b) Odorant stimulation promotes survival of rodent olfactory receptor neurons via PI3K/Akt activation and Bcl-2 expression. Mol Cells 38:535–539.

Kurahashi T, Lowe G, Gold, GH. (1994) Suppression of odorant responses by odorants in olfactory receptor cells. Science 265:118–120.

Laing DG, Panhuber H, Wilcox ME, Pittmann EA (1984) Quality and intensity of binary odor mixtures. Physiol. Behav. 33:309–319.

Laing DG, Wilcox ME (1987) An investigation of the mechanism of odor suppression using physical and dichorhinic mixtures. Behav. Brain Res. 26:79–87.

Latorraca NR, Venkatakrishnan AJ, Dror RO (2016) GPCR dynamics: Structures in motion. Chem Rev:acs.chemrev.6b00177.

Lohse MJ, Hofmann KP (2015) Spatial and temporal aspects of signaling by G protein– coupled receptors. Mol Pharmacol 88.

Lowe G, Gold GH (1991) The spatial distribution of odorant sensitivity and odor-induced currents in salamander olfactory receptor cells. J Gen Physiol 442:147–68.

Luttrell LM (2014) Minireview: More than just a hammer: ligand “bias” and pharmaceutical discovery. Mol Endocrinol 28:281–294.

Manglik A, Kim TH, Masureel M, Altenbach C, Yang Z, Hilger D, Lerch MT, Kobilka TS, Thian FS, Hubbell WL, Prosser RS, Kobilka BK (2015) Structural insights into the dynamic process of β2-adrenergic receptor signaling. Cell 161:1101–1111.

Mashuo I, Ostrovkaya O, Kramer GM, Jones CD, Xie K, Martemyanov KA (2015) Distinct profiles of functional discrimination among G proteins determine the actions of G protein-coupled receptors. Science Signaling 8:1–16.

Mashukova A, Spehr M, Hatt H, Neuhaus EM (2006) Beta-arrestin2-mediated internalization of mammalian odorant receptors. J Neurosci 26:9902–9912.

Mayer U, Küller A, Daiber PC, NeudorfI, Warnken U, Schnölzer M, Frings S, Möhrlen F (2009) The proteome of rat olfactory sensory cilia. Proteomics 9:322–334.

McEwen DP, Jenkins PM, Martens JR (2008) Olfactory cilia: our direct neuronal connection to the external world. Curr Top Dev Biol 85:333–370.

McIntyre JC, Joiner AM, Zhang L, Iñiguez-Lluhí J, Martens JR (2015) SUMOylation regulates ciliary localization of olfactory signaling proteins. J Cell Sci 128:1934–1945.

Munger SD (2009) Olfaction: Noses within noses. Nature 459:521–522.

Nickell MD, Breheny P, Stromberg AJ, McClintock TS (2012) Genomics of mature and imature olfactory sensory neurons. J Comp Neurol 520:2608–2629.

Oboti L, Pérez-Gómez A, Keller M, Jacobi E, Birnbaumer L, Leinders-Zufall T, Zufall F, Chamero P (2014) A wide range of pheromone-stimulated sexual and reproductive behaviors in female mice depend on G protein Gαo. BMC Biol 12:31.

Oka Y, Omura M, Kataoka H, Touhara K (2004) Olfactory receptor antagonism between odorants. Embo J 23:120–126.

Omura M, Mombaerts, P (2014) Trpc2-expressing sensory neurons in the ain olfactory epitheium. Cell Reports 8:583–595.

Orr AW, Pallero MA, Murphy-Ullrich JE (2002) Thrombospondin stimulates focal adhesion disassembly through Gi- and phosphoinositide 3-kinase-dependent ERK activation. J Biol Chem 277:20453–20460.

Rameh LE, Cantley LC (1999) The role of phosphoinositide 3-kinase lipid products in cell function. J. Biol Chem 274:8347–8350.

Reddy G, Zak J, Vergassola M, Murthy VN (2018) Antagonism in olfactory receptor neurons and its implications for the perception of odor mixtures. eLife 2018:7:e34598. DOI: 10.7554/eLife.34958.

Rospars JP, Lansky P, Chaput M, Duchamp-Viret P (2008) Competitive and noncompetitive odorant interactions in the early neural coding of odor mixtures. J Neurosci 28:2659–66.

Sanz G (2005) Comparison of odorant specificity of two human olfactory receptors from different phylogenetic classes and evidence for antagonism. Chem Senses 30:69–80.

Saraiva LR, Ibera-Soria X, Kahn M, Omura M, Scialdone A, Mombaerts P, Marioni JC, Logan SW (2015) Hierarchial deconstruction of mouse olfactory sensory neurons: from whole mucosa to single-cell RNA-seq. Sci Reports 5:18178.

Schandar M, Laugwitz KL, Boekhoff I, Kroner C, Gudermann T, Schultz G, Breer H (1998) Odorants selectively activate distinct G protein subtypes in olfactory cilia. J Biol Chem 273:16669–16677.

Scholz P, Kalbe B, Jansen F, Altmueller J, Becker C, Mohrhardt J, Schreiner B, Gisselmann G, Hatt H, Osterloh S (2016a) Transcriptome analysis of murine olfactory sensory neurons during development using single cell RNA-Seq. Chem Senses 313–323.

Scholz P, Mohrhardt J, Jansen F, Kalbe B, Haering C, Klasen K, Hatt H, Osterloh S (2016b) Identification of a novel Gnao-mediated alternate olfactory signaling pathway in murine OSNs. Front Cell Neurosci 10:63.

Shepard BD, Natarajan N, Protzko RJ, Acres OW, Pluznick JL (2013) A cleavable N-terminal signal peptide promotes widespread olfactory receptor surface expression in HEK293T cells. PLoS One 8:e68758.

Shirokova E, Schmiedeberg K, Bedner P, Niessen H, Willecke K, Raguse J-D, Meyerhof W, Krautwurst D (2005) Identification of specific ligands for orphan olfactory receptors. G protein-dependent agonism and antagonism of odorants. J Biol Chem 280:11807–11815.

Slepak VZ, Wilkie TM, Simon MI (1993) Mutational analysis of G protein alpha subunit G(o) alpha expressed in *Escherichia coli*. J Biol Chem 268:1414–1423.

Smrcka A V. (2015) Fingerprinting G protein-coupled receptor signaling. Sci Signal 8:fs20–fs20.

Spehr M, Wetzel CH, Hatt H, Ache BW (2002) 3-phosphoinosiides modulate cyclic nucleotide signaling in olfactory receptor neurons. Neuron 33:731–739.

Steiner D, Saya D, Schallmach E, Simonds WF, Vogel Z (2006) Adenylyl cyclase type-VIII activity is regulated by Gβγ subunits. Cell Signal 18:62–68.

Su C-Y, Menuz K, Reisert J, Carlson JR (2012) Non-synaptic inhibition between grouped neurons in an olfactory circuit. Nature 492: 66–72.

Takeuchi H, Ishida H, Hikichi S, Kurahashi T (2009) Mechanism of olfactory masking in the sensory cilia. J. Gen Physiol. 133:583–601.

Touhara K, Sengoku S, Inaki K, Tsuboi A, Hirono J, Sato T, Sakano H, Haga T (1999) Functional identification and reconstitution of an odorant receptor in single olfactory neurons. Proc Natl Acad Sci U S A 96:4040–4045.

Tsuboi A, Imai T, Kato HK, Matsumoto H, Igarashi KM, Suzuki M, Mori K, Sakano H (2011) Two highly homologous mouse odorant receptors encoded by tandemly-linked MOR29A and MOR29B genes respond differently to phenyl ethers. Eur J Neurosci 33:205–213.

Ukhanov K, Brunert D, Corey EA, Ache BW (2011) Phosphoinositide 3-kinase-dependent antagonism in mammalian olfactory receptor neurons. J Neurosci 31:273–280.

Ukhanov K, Corey EA, Ache BW (2013) Phosphoinositide 3-kinase dependent inhibition as a broad basis for opponent coding in mammalian olfactory receptor neurons. PLoS One 8:e61553.

Ukhanov K, Corey EA, Brunert D, Klasen K, Ache BW (2010) Inhibitory odorant signaling in mammalian olfactory receptor neurons. J Neurophysiol 103:1114–1122.

Venkatakrishnan AJ, Deupi X, Lebon G, Heydenreich FM, Flock T, Miljus T, Balaji S, Bouvier M, Veprintsev DB, Tate CG, Schertler GFX, Babu MM (2016) Diverse activation pathways in class A GPCRs converge near the G-protein-coupling region. Nature 536:484–487.

Wang Q, Titlow WB, McClintock DA, Stromberg AJ, McClintock TS (2017) Activity-dependent gene expression in the mammalian olfactory epithelium. Chem Senses 42:611–624.

Watt WC, Sakano H, Lee Z-Y, Reusch JE, Trinh K, Storm DR (2004) Odorant stimulation enhances survival of olfactory sensory neurons via MAPK and CREB. Neuron 41:955–967.

Wekesa KS, Anholt RR (1999) Differential expression of G proteins in the mouse olfactory system. Brain Res 837:117–126.

Wettschureck N (2005) Mammalian G proteins and their cell type specific functions. Physiol Rev 85:1159–1204.

Wingler LM, Elgeti M, Hilger D, Latorraca NR, Lerch MT, Stus DP, Dror RO, Koblika BK, Hubbell, WL, Lefkowitz RJ (2019) Angiotensin analogs with divergent bias stabilize distinct receptor conformations. Cell 176:468–478.

Xu L, Wenze L, Venkatakaushik V, Hillman EMC, Firestein S (2019) Widespread receptor driven modulation in peripheral olfactory coding. bioRxiv Sept 8, 2019: doi:http://dx.doi.org/10.1101/760330.

Yu Y, Boyer NP, Zhang C (2014) Three structurally similar odorants trigger distinct signaling pathways in a mouse olfactory neuron. Neuroscience 275:194–210.

Zhainazarov AB, Spehr M, Wetzel CH, Hatt H, Ache BW (2004) Modulation of the olfactory CNG channel by Ptdlns(3,4,5)P3. J Memb Biol 201:51–57.

Zhang G, Ttilow WB, Biecker SM, Stromberg AJ, McClintock TS (2016) Lhx2 determines odorant receptor expression frequency in mature olfactory sensory neurons, eNeuro 3(5) pii ENEURO 0230-16.2016.

Zhuang H, Matsunami H (2007) Synergism of accessory factors in functional expression of mammalian odorant receptors. J Biol Chem 282:15284–15293.

Zolotukhin S, Potter M, Zolotukhin I, Sakai Y, Loiler S, Fraites TJ, Chiodo VA, Phillipsberg T, Muzyczka N, Hauswirth WW, Flotte TR, Byrne BJ, Snyder RO (2002) Production and purification of serotype 1, 2, and 5 recombinant adeno-associated viral vectors. Methods 28:158–167.

